# Chromatin condensation delays human mesenchymal stem cells senescence by safeguarding nuclear damages during long term *in vitro* expansion

**DOI:** 10.1101/2023.04.22.537784

**Authors:** Rohit Joshi, Tejas Suryawanshi, Sourav Mukherjee, Shobha Shukla, Abhijit Majumder

**Affiliations:** Department of Chemical Engineering, Indian Institute of Technology, Bombay, India, 400076; Centre for research in Nano Technology and Science, Indian Institute of Technology, Bombay, India, 400076; Department of Metallurgical Engineering and Materials Science, Indian Institute of Technology, Bombay, India, 400076

## Abstract

Human mesenchymal stem cells (hMSCs) are multipotent cells that can differentiate into adipocytes, chondrocytes and osteoblasts. Due to their differentiation potential, hMSCs are among the most frequently used cells for therapeutic applications in tissue engineering and regenerative medicine. However, the number of cells obtained through isolation alone is insufficient for hMSC-based therapies and basic research, necessitating their *in-vitro* expansion. Conventionally, this is often carried out on rigid surfaces such as tissue culture petriplates (TCPs). However, during *in-vitro* expansion, hMSCs lose their proliferative ability and multilineage differentiation potential, making them unsuitable for clinical use. Although multiple approaches have been tried to maintain hMSC stemness over prolonged expansion, finding a suitable culture system to achieve this remains an unmet need. Recently, few research groups including ours have shown that hMSCs maintain their stemness over long passages when cultured on soft substrate. In addition, it has been shown that hMSCs cultured on soft substrates have more condensed chromatin and lower levels of histone acetylation compared to those cultured on stiff substrates. It has also been shown that condensing/decondensing chromatin by deacetylation/acetylation can delay/hasten replicative senescence in hMSCs during long-term expansion on TCPs. However, how chromatin condensation/decondensation influences nuclear morphology and DNA damage - which are strongly related to the onset of senescence and cancer - is still not known.

To answer this question, here we cultured hMSCs for long duration (P4-P11) in presence of epigenetic modifiers histone acetyltransferase inhibitor (HATi) which promotes chromatin condensation by preventing histone acetylation and histone deacetylase inhibitor (HDACi) which promotes chromatin decondensation and investigated their effect on various nuclear markers related to senescence and cancer. We have found that consistent acetylation causes severe nuclear abnormalities whereas chromatin condensation by deacetylation helps in safeguarding nucleus from damages caused by *in-vitro* expansion.

## Introduction

Human mesenchymal stem cells (hMSCs) are multipotent stem cells that have ability to differentiate into variety of cell type including adipocytes, chondrocytes, osteoblast and myoblast (Engler et al., 2006; Pittenger et al., 1999). In addition, due to their paracrine activity hMSCs can secrete a variety of cytokines and chemokines having immunomodulatory effect (Cheleuitte et al., 1998; Kyurkchiev, 2014; Madrigal et al., 2014). Owning to these excellent properties hMSCs are ideal cell source for tissue engineering and regenerative medicine (TERM) and most often used stem cells in clinical trials. Although the number of clinical studies has increased by three times over the last decade, the proportion of studies reaching to phase IV trails are very few (Bunpetch et al., 2019; Trounson & McDonald, 2015). While there are several interrelated issues that contribute to the paucity of late phase trials, one issue is the lack of effective method for *in-vitro* expansion of hMSCs which retain the differentiation and therapeutic qualities. As the number of cells obtained through isolation alone is insufficient for basic research and/or therapeutic applications, so hMSCs are expanded *in-vitro*. In traditional scientific approach this is often carried out by culturing hMSCs on rigid surfaces such as tissue culture petri plastics (TCPs). As for clinical applications, millions of cells per kg patient body weight is required therefore, the successful *in-vitro* expansion of hMSCs is a prerequisite for their clinical applications (Ren et al., 2012; Squillaro et al., 2016). However, during *in-vitro* expansion when hMSCs were expanded on TCPs they enter in senescence state and decreases their differentiation, secretory potential, migration and homing ability making them unsuitable for clinical use (Bonab et al., 2006; De Becker & Van Riet, 2016; Wagner et al., 2010). Like any other primary somatic cells, after few passages, hMSCs undergoes progressive decline in fitness as they enter into senescence state, which is morphologically characterized by enhanced spreading area and shape irregularity (Zaim et al., 2012). Thus, there is a need for culture strategies that can maintain hMSCs fitness and potency during long term expansion. Though multiple approaches have been tried to maintain MSC stemness over prolonged expansion, finding an easy-to-use culture system to achieve the same is still an unmet need (Saei Arezoumand et al., 2017). Recently, few research groups including our have shown that hMSCs maintain their stemness over long passages when cultured on an optimally soft polyacrylamide (PAA) gel maintaining their cellular morphology and proliferative potential with delay in senescence (Kureel et al., 2019a; Rao et al., 2019). Previous studies have also shown that mechanical cues such as substrate stiffness influences chromatin remodelling and epigenome of hMSCs (Killaars, Grim, Walker, Hushka, Brown, Anseth, et al., 2019; Miroshnikova et al., 2017). Chromatin remodelling primarily regulates gene expression through epigenetic modifications such as acetylation, methylation, and phosphorylation. The acetylation landscape is highly dynamic and governed by two classes of enzymes—histone acetyltransferases (HATs) and histone deacetylases (HDACs). Acetylation of histones by HATs leads to chromatin decondensation enabling gene expression. Alternatively, deacetylation of histone by HDACs results in condensation of chromatin and repression of gene expression. Recently it has been shown that hMSCs cultured on stiff substrate have more decondensed chromatin with higher histone acetylation as compared to nuclei on stiff substrate (Joshi et al., 2022; Killaars, Grim, Walker, Hushka, Brown, Anseth, et al., 2019).

As discussed previously it is now known that replicative senescence in hMSCs can be delayed by culturing them on soft substrate and soft substrate promote chromatin condensation. Motivated by these findings we hypothesized that changing epigenetic landscape (acetylation/deacetylation) of hMSCs during long term expansion play role in regulating senescence. So, we sought to investigate how culturing hMSCs on TCPs for long term expansion in the presence of epigenetic modifiers HATi (Promotes chromatin condensation) and HDACi (promotes chromatin decondensation) effects hMSCs senescence. We cultured hMSCs in presence of anacardic acid (HATi) and valproic acid (HDACi) during serial passing (P4-P11) on TCPS. Our results suggest that culturing hMSCs in the presence of anacardic acid delays cellular senescence by safeguarding nuclear morphology as compared for hMSCs cultured in the presence of valproic acid and control. Results from these studies will help the scientific community to identify the chromatin based critical parameters for expansion of hMSCs with highly therapeutic and regenerative potency besides material-based strategies.

## Material and methods

### Cell culture

Bone marrow-derived hMSCs were purchased from Lonza (Cat.No. #PT-2501). hMSCs were cultured in Low glucose DMEM (Himedia; AL006) supplemented with 16% FBS (Himedia; RM9955), 1% Antibacterial-Antimycotic (Himedia; A002) and 1% Glutamax (Gibco; 35050) under humidified conditions at 37°C with 5% CO2. The cells were trypsinized with TrypLE™(Gibco; 12604021) once they reached the confluency of 70%.

### Treatment with HDACi and HATi

To check the effect of HDACi and HATi, 0.5mM of Valproic acid (PHR 1061) and 30µM of anacardic acid (ANA) was added to the growth media after 24hrs of seeding. For differentiation experiments, VA and ANA was added with differentiation media.

### Differentiation assays

hMSCs were seeded at 2000 cells/cm^2^ in a 12-well culture plate in growth medium for 24h followed by differentiation media. Adipogenic (Invitrogen, A10410) and osteogenic (Invitrogen, A10069) differentiation kits were used. Cells were incubated for 9 days in adipo and 14 days for osteo induction media before quantitative assays. Differentiation media change was given after every third day. After the completion of differentiation duration, cells were fixed with 4% paraformaldehyde (PFA) for 30min at room temperature followed by staining with Oil Red O (SigmaO0625) for adipogenic differentiation and Alizarin Red (Sigma A5533) for osteogenic differentiation. After incubating with staining solution for 20 min samples were washed thrice with DPBS (adipo) or MilliQ (osteo). Images were captured for quantitative analysis using EVOS inverted microscope (Invitrogen) in bright-field colour channel.

### Immunofluorescence staining

Cells were fixed with 4% PFA in PBS for 15 min at room temperature (RT) then washed with PBS thrice. Cells are then permeabilized with permeabilizing buffer (0.5% Triton X-100-Sigma Aldrich in CSB) for 10 min and blocked with BSA (4% Bovine serum albumin in PBS) for 30 min to minimize nonspecific protein binding. Anti-lamin A (1:500, mouse, Abcam, Cat. No. 8980), anti-AcK (1:500, rabbit, Abcam, Cat. No. 190479) primary antibodies in 4% BSA were added to the samples and incubated overnight at 4°C. Primary antibodies were removed and samples were rinsed with PBS two times for 10 min. Samples were then incubated at room temperature with secondary antibodies (1:500, donkey anti-mouse AlexaFlour 488 Cat. No. 21202), donkey anti rabbit AlexaFlour 569 Cat. No. ab175470) phalloidin (1:400, AlexaFlour 532, Cat. No. A22282) with Hoechst 33342 (Cat. No. H3570) of dilution 1:5000 in 4% BSA for 2 hrs at room temperature. Thereafter the samples were rinsed twice with PBS. All immunostained samples were stored in PBS at 4°C until Imaging. All samples are imaged at 63X (oil) magnification using laser scanning Confocal Microscope (LSM, Carl Zeiss).

### BrdU assay

To check the percentage of S-phase cells in the cell cycle, cells from EP (early passage) and LP (late passage) were trypsinized and seeded on glass coverslips. After 48 hrs of seeding, BrdU reagent (Invitrogen, 000103) was added in 1:100 (v/v) ratio in media and incubated for 4 hrs at 37°C in a humidified incubator with 5% CO_2_. Thereafter cells were fixed (4% paraformaldehyde), permeabilized (0.5% Triton-X), denatured (2 M HCL), blocked (1.5% bovine serum albumin), and incubated with anti-BrdU antibody (Invitrogen, B35128, 1:100) and counterstained with AlexaFluor 568 (Invitrogen, A11061, 1:400). Immunofluorescence images were captured using EVOS-FL auto and BrdU positive and negative cells were counted manually using ImageJ.

### Senescence assays

Senescence-associated β-galactosidase (SA-β-gal) was used to detect MSCs senescence using SA-β-gal staining kit (Abcam, AB65351) according to the manufacturer’s instructions. Briefly, cells from EP and LP were seeded in a six-well plate and incubated in growth media. Afterwards, cells were fixed, stained with β-gal solution and incubated at 37°C without CO_2_. 10–15 random images were captured for each condition for analysis and Images were captured using EVOS inverted microscope (Invitrogen) in bright-field colour channel.

## Results

### HATi increases chromatin condensation of hMSCs cultured during long term expansion

Epigenetic modifications of histones are associated with senescence in different species and cells (Crouch et al., 2022). Histone acetylation is one of such critical epigenetic modifications. Here, we sought to investigate how a change in global acetylation state of histones may influence the replicative senescence in long term *in-vitro* expansion of hMSCs. Towards that goal, we cultured hMSCs for multiple passages (P4-P11) in presence of valproic acid (VA) and anacardic acid (ANA), two well established inhibitors of histone acetylation (HDACi) and histone deacetylation (HATi) respectively (Fig. 1). While HDACi keeps the chromatin in an open state by hyperacetylating the histones (Fig. 1B), HATi keeps it in a more condensed state by inhibiting the acetylation process (Fig. 1C). Our results showed that for early passage with respect to (normal media) NM (Fig. 2A), there was a significant decrease in global histone acetylation in hMSCs cultured in the presence of ANA (∼25%) (Fig. 2B and G) and a significant increase in the presence of VA (∼1.7 times) (Fig. 2C and G). Furthermore, when we cultured hMSCs for the long term from P4 to P11, we found that in the late passage, there was no significant difference in histone acetylation for NM (Fig. 2D and G) as compared to early passage. When cultured in the presence of inhibitors HATi and HDACi, we found that in late passage, there was a significant decrease in histone acetylation in the presence of ANA (∼45%) (Fig. 2E & G), and no difference when cultured with VA (Fig. 2F & G) as compared to early passage.

**Figure 1.**
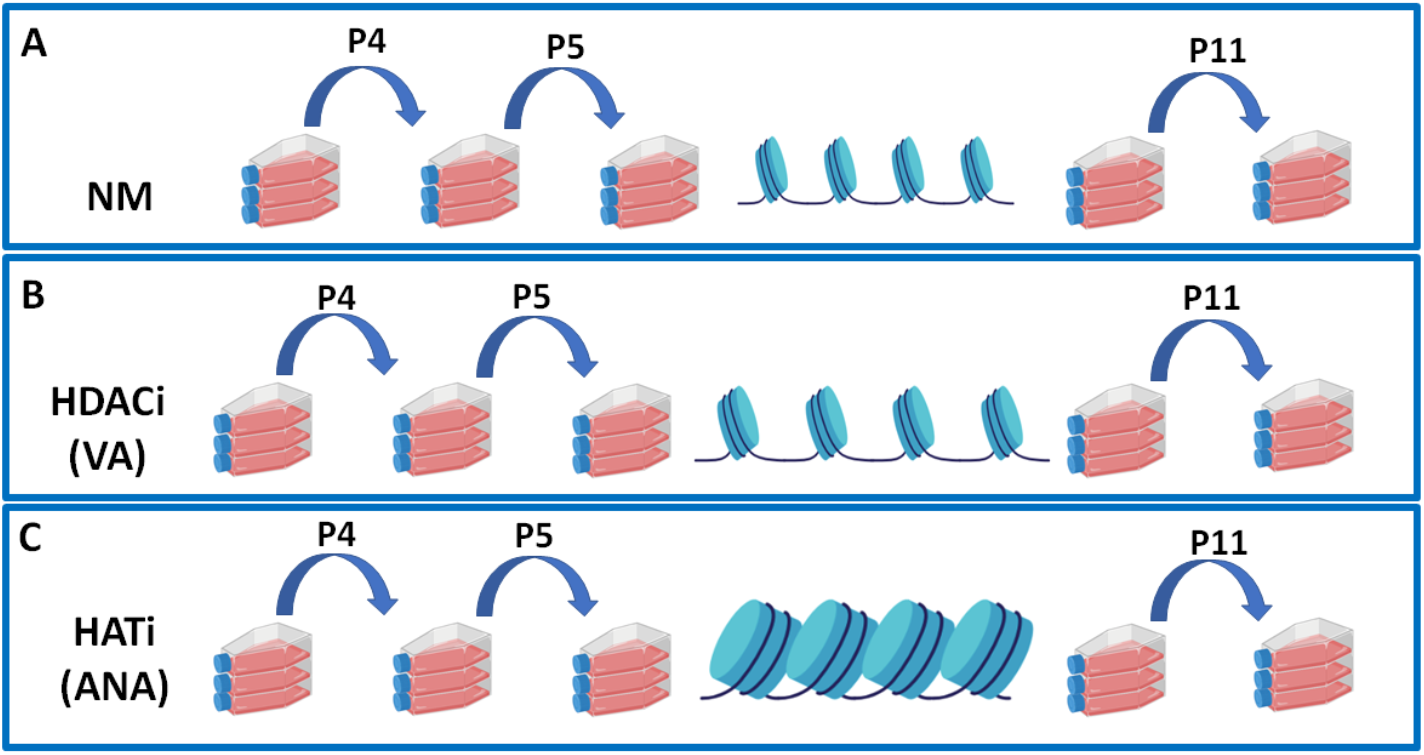
Schematic diagram showing the Experimental workflow: Serial passaging of hMSCs was done in TCPs from P4 to P11 cultured in A) Control (NM) B) in presence of VA C) in presence of ANA.

**Figure 2.**
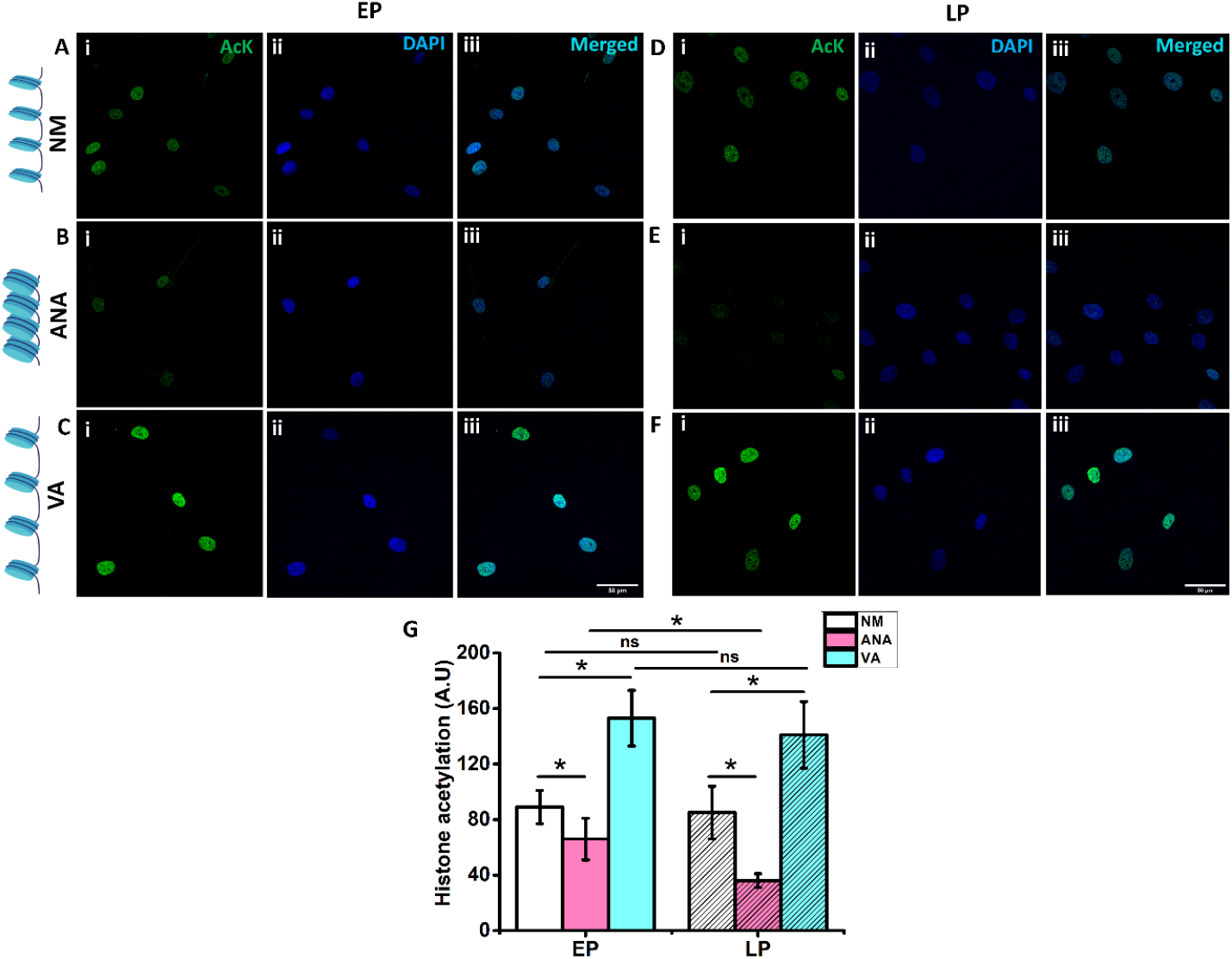
Staining of (i) global histone acetylation (Ack) (ii) DAPI, and (iii) merged image of nuclei when the cells were in (A; EP and D; LP) control (NM, without any inhibitor), (B; EP and E; LP) growth media with anacardic acid (HATi) and (C; EP and F; LP) growth media with valproic acid (HDACi). Blue; DAPI, Green; AcK, Cyan; merged. (G) Graph comparing histone acetylation under various conditions (NM-normal media, ANA-anacardic acid, VA – valproic acid, n> 60 nuclei, Scale bar = 50 µm *p<0.05, ns = non-significant).

### Chromatin condensation helps in maintaining hMSCs proliferation during long term expansion

Next, we investigated the effect of chromatin condensation and decondensation on hMSCs proliferation. BrdU incorporation assay was used to identify the proliferating cells. Our results show that in early passage there was no significant difference in hMSCs proliferation between NM (Fig. 3A and G), and ANA (Fig. 3B and G), but a significant decrease was observed in the presence of VA (∼24%) (Fig. 3C and G) as compared to NM, showing that hyperacetylation of histones by HDACi (VA) negatively influences the cell cycle. When hMSCs were cultured on TCPs for long term there was significant decrease in proliferation in all three cases in LP as compared to EP. We found that in late passage, hMSCs cultured in the presence of NM showed a significant decrease in proliferation (∼55%) (Fig. 3D and G) as compared to early passage. When cultured in the presence of inhibitors HATi and HDACi, ANA showed a minimum decrease in proliferation (∼33%) (Fig. 3E and G), whereas VA showed a maximum decrease in proliferation (∼70%) (Fig. 3F and G) in LP as compared to EP. Moreover, we also checked for the cumulative population doubling (CPD) for hMSCs cultured in all three conditions and found that cells cultured in the presence of ANA grows almost linearly up to P9 while CPD declined drastically for cells cultured in VA from P6 (Fig. 3H) resulting into a difference in population doubling between ANA and VA at P10 to be ∼8. In other words, culturing the hMSCs in presence of ANA would result into 2^8^ (256) times more cells than the same cultured in VA.

**Figure 3.**
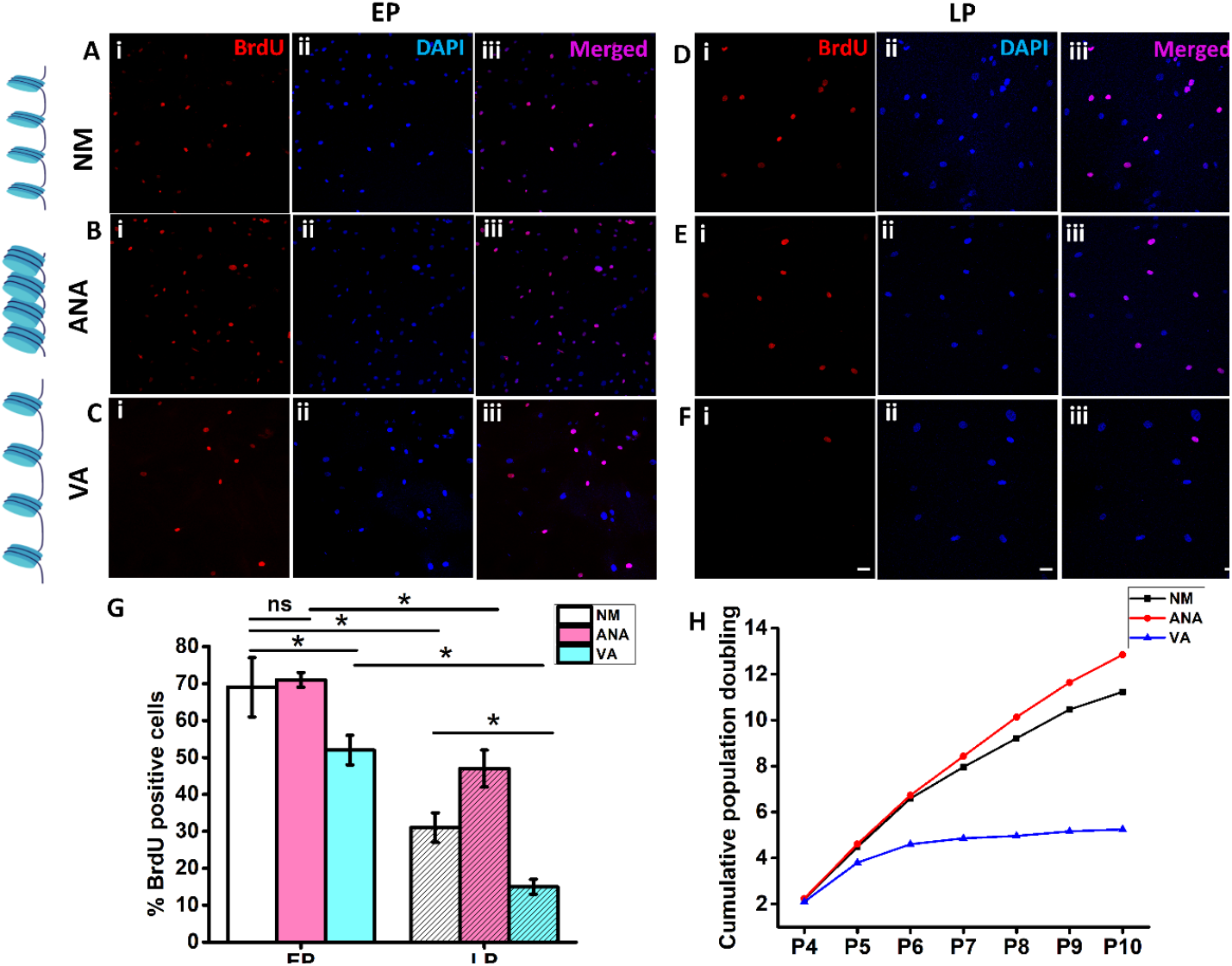
BrdU incorporation study to estimate proliferation of early passage (EP, A-C) & late passage (LP, D-F) for hMSCs cultured in normal media (NM, A and D), with anacardic acid (ANA, B and E), and with valproic acid (VA, C and F). Blue; DAPI, Red; Nuclei with BrdU, Magenta; merged. (G) Graph comparing BrDU incorporation under various conditions; (H) graph shows cumulative population doubling of hMSCs cultured under all three conditions (n> 400 nuclei, Scale bar= 50 µm, *p<0.05, ns = non-significant).

### Chromatin compaction delays hMSCs senescence

To check the effect of HDACi and HATi on hMSCs senescence, we checked for β-gal activity, which is the gold standard assay to check senescence. Our results showed that there was no significant difference in β-gal activity for all three conditions for hMSCs (NM, ANA, VA) cultured during early passage (P4), as shown in (Fig. 4A-C and G). Whereas, when we checked for late passage (P11), we found that there was a significant increase in β-gal activity in all three conditions, as shown in (Fig. 4D-F and G). When cultured in NM, a significant increase (∼3.3 times) (Fig. 4D and G) in LP as compared to EP. When cultured in the presence of HATi and HDACi, cells cultured with ANA showed a minimum increase (∼1.5 times) (Fig. 4E and G), and VA showed a maximum increase (∼ 5 times) (Fig. 4F and G) in β-gal activity in LP compared to EP. In conclusion our results showed that culturing hMSCs for long term expansion in condensed chromatin state in the presence of HATi delays hMSCs senescence.

**Figure 4.**
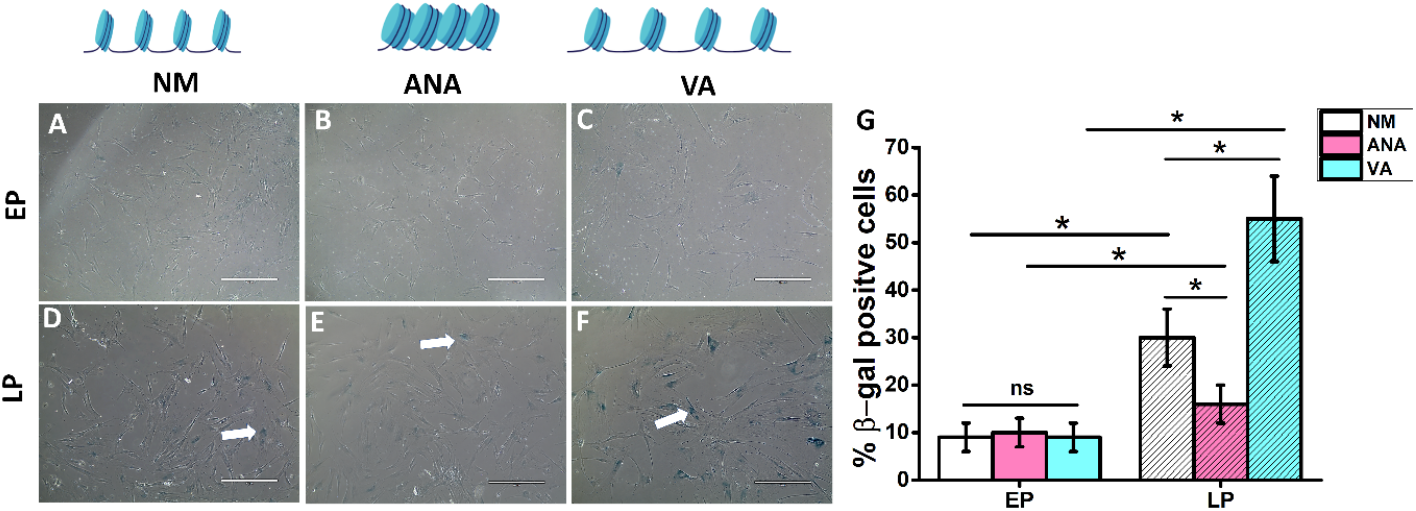
Investigation of senescence in early passage (EP) (A-C) and late passage (LP) (D-F) in presence of normal media (NM, A and D) with anacardic acid (ANA, B and E), and with valproic acid (VA, C and F). (G) Percentage of β-gal positive cells in various conditions (NM, ANA, VA) showing highest β-gal activity for hMSCs cultured in presence of VA lowest in presence of ANA for late passage cells (Scale bar = 400 µm, *p<0.05, ns = non-significant).

### Chromatin condensation maintains adipogenic differentiation potential during long expansion

Previous studies have shown that hMSCs differentiation ability is passage dependent and delays with senescence (Turinetto et al., 2016). Motivated by these findings we wanted to check the effect of chromatin condensation/decondensation on the adipogenic differentiation potential of hMSCs cultured during serial passaging. Our results showed that with respect to NM (Fig. 5A), there was a significant increase in the percentage of ORO-positive cells in the ANA (∼1.3 times) (Fig. 5B & G) and a significant reduction when cultured in the presence of VA (∼43%) (Fig. 5C, & G).

**Figure 5.**
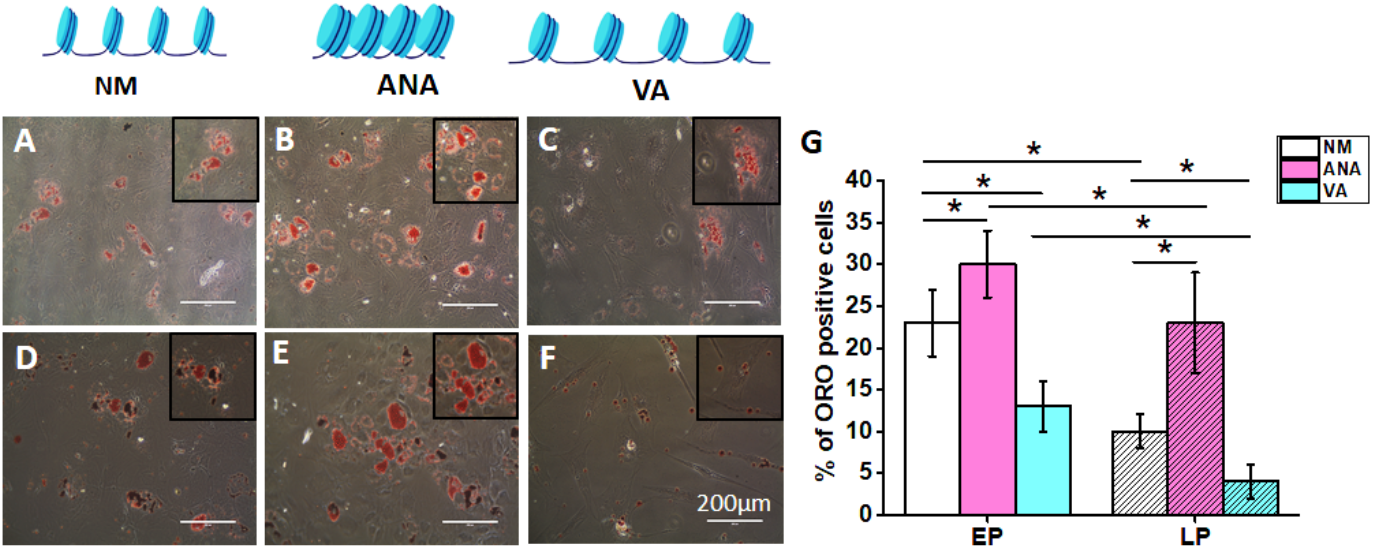
Investigation of adipogenic differentiation of hMSCs by oil red o staining on early passage hMSCs in (A) ctrl (NM), (B) presence of anacardic acid and (C) presence of valproic acid and in late passage hMSCs in (D) ctrl (NM), (E) presence of anacardic acid and (F) presence of valproic acid; (G) Graph comparing percentage of oil red o positive cells under various conditions (NM-normal media, ANA-anacardic acid, VA – valproic acid); (Scale bar = 200 µm, *p<0.05).

Furthermore, when we cultured hMSCs for long term there was decline in adipogenic differentiation potential in all three cases in LP as compared to EP (Fig. 5G). We found that in late passage, hMSCs cultured in NM showed a significant decrease (∼56%) in adipogenic differentiation potential (Fig. 5D and G) compared with the late passage. When cultured in the presence of HATi and HDACi, cells cultured in the presence of ANA showed a minimum decline (∼23%) (Fig. 5E and G), and VA showed a maximum decline (∼69%) (Fig. 5F & G) in LP as compared to EP.

Supporting our previous results hMSCs cultured during long term in presence of ANA delays hMSCs senescence while maintaining adipogenic differentiation potential as compared to NM and VA. Overall, our findings demonstrate that culturing hMSCs during serial passaging in the presence of ANA decreases hMSCs senescence and rescues adipogenic differentiation potential.

### Chromatin condensation safeguards nuclear morphology during long term expansion

It is now known that distorted nuclear morphology including abnormalities such as nuclear blebbing are strongly associated with cellular senescence (Heckenbach et al., 2022; Pathak et al., 2021). Senescent cells show prominent nuclear blebs that change the overall 3D morphology of the nuclei. In the previous section, we demonstrated that cells cultured for multiple passages in the presence of ANA (deacetylation) showed less β-gal activity, which is a standard marker for checking cellular senescence. However, the influence of persistent acetylation/deacetylation on nuclear morphometric parameters during long-term expansion remains unexplored. To fill this gap, we investigated the effect of chromatin condensation/decondensation on nuclear morphometric parameters in hMSCs cultured on TCPs during serial passaging. We compared the early and late passage nuclear morphology, bleb and nuclear abnormalities for hMSCs cultured in all three conditions (NM, ANA, VA). First, we analyzed the nuclear morphological parameters (projected area, volume, surface area, and circularity) and found that in early passages, there was no significant difference between NM and ANA for any of the nuclear morphometric parameters (Fig. 6A, B, G, and H). However, there was a significant increase in the nuclear morphometric parameters for cells cultured in the presence of VA (area∼ 40%, volume∼34%, surface area∼34%) compared to NM (Fig. 6A, C, G and H). Furthermore, when hMSCs were cultured for long term there were significant differences in all the nuclear morphometric parameters for all three conditions in LP as compared to EP. During late passage, when cultured in the presence of NM, there was a significant increase in the nuclear projected area (∼59%, Fig. 6D and G), volume (∼67%), and surface area (∼61%) as compared to early passage. However, when cultured in the presence of HATi and HDACi, we found that hMSCs cultured in the presence of ANA showed a minimum increase in all nuclear morphometric parameters, nuclear projected area (∼15%, Fig. 6G) and nuclear volume (∼44%; Fig. 6H), surface area (∼37%, Fig. S1) and no significant change in circularity (Fig. S1) in LP as compared to EP. When cultured in the presence of VA, the highest increase was observed in the projected nuclear area (∼72%, Fig. 6G), volume (∼70%, Fig. 6H), surface area (∼60%, Fig. S1) and significant decrease in circularity (∼6%, Fig. S1) in LP as compared to EP. Furthermore, we checked the effect of HDACi and HATi on nuclear blebbing and abnormalities. For the early passage, there was no significant difference in nuclear abnormalities between all three conditions (Fig. 6I) and no significant difference in nuclear blebbing between NM and ANA but VA shows ∼80% increase in nuclear blebbing as compared to NM (Fig. 6J). Furthermore, when hMSCs were cultured for a long period, there was a significant increase in nuclear abnormalities (except ANA) and nuclear blebbing under all three conditions in LP compared to EP (Fig. 6A-F, I and J). We showed that in late passage cells cultured in NM shows significant increase in nuclear abnormalities (∼32%) and blebbing (∼260%) (Fig. 6I and J) compared to that in the early passage. When cultured in the presence of inhibitors HATi and HDACi, we showed that hMSCs cultured in the presence of ANA showed a minimum increase in nuclear abnormalities (∼11%) and nuclear blebs (∼133%) and maximum increases in nuclear abnormalities (∼97%) and blebs (∼377%, Fig. 6F and J) when cultured in the presence of VA in LP compared to EP. Overall, our data suggest that long-term culture of hMSCs in the presence of HATi (ANA) prevents deterioration of nuclear morphology, nuclear blebs, and abnormalities, which are markers of cellular senescence and aging.

**Figure 6.**
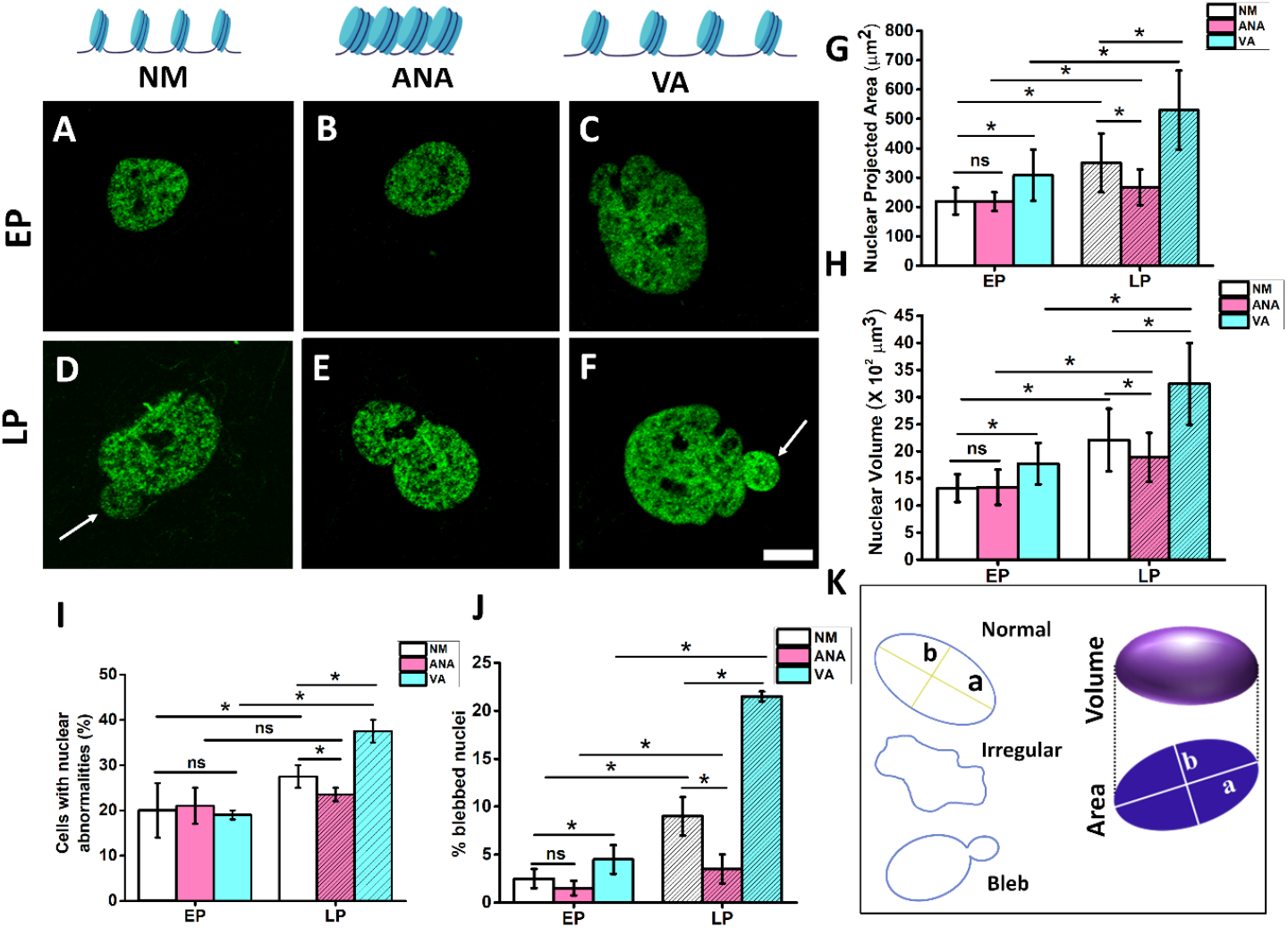
Chromatin condensation safeguards nuclear morphometric parameters during long term expansion. Immunofluorescence images of acetylation showing nuclear blebbing in early passage (EP, A-C) & late passage (LP, D-F) for hMSCs cultured in normal media (NM, A and D), with anacardic acid (ANA, B and E), and with valproic acid (VA, C and F). Graph comparing (G) change in nuclear projected area, (H) volume, (I) percentage of cells with nuclear abnormalities and (J) percentage of cells with nuclear bleb (K) Schematic showing abstract representation of normal, irregular and blebbed nucleus (n> 65 nuclei, Scale bar = 10µm *p<0.05, ns = non-significant).

### Lamin A expression increases with serial passaging

Expression levels of lamins are one of the important senescence markers as previously people have shown there is an accumulation of lamin A and loss of lamin B in senescent cells. Therefore, we investigated the effects of HDACi (chromatin decondensation) and HATi (chromatin condensation) on Lamin A expression during serial passaging. We observed that hMSCs cultured in early passage (P4) had no significant difference in total Lamin A intensity under all three conditions (NM, ANA, and VA) (Fig. 7A-C & G). Moreover, we found that in early passage cells, Lamin A was more localized to the nuclear periphery as compared to the nucleoplasm as shown in (Fig. 7A-C). Next, we investigated lamin A expression in late passage cells in the presence of all three conditions and found that there was a significant increase in lamin A expression in all three cases (NM, VA, and ANA) compared to early passage cells. We found that late-passage cells cultured in NM showed a significant increase in Lamin A intensity (∼1.5 times) (Fig. 7D and G) compared to early passage cells. When cultured in the presence of ANA and VA, cells in ANA showed a minimum increase (∼1.5 times) (Fig. 7E and G) and VA showed the maximum increase (∼1.9 times) in Lamin A expression in LP with respect to EP. From our results, we also found that in late passage cells lamin A was not only localized in the nuclear periphery but also accumulated in the nucleoplasm, as shown in (Fig. 7H, I &J).

**Figure 7.**
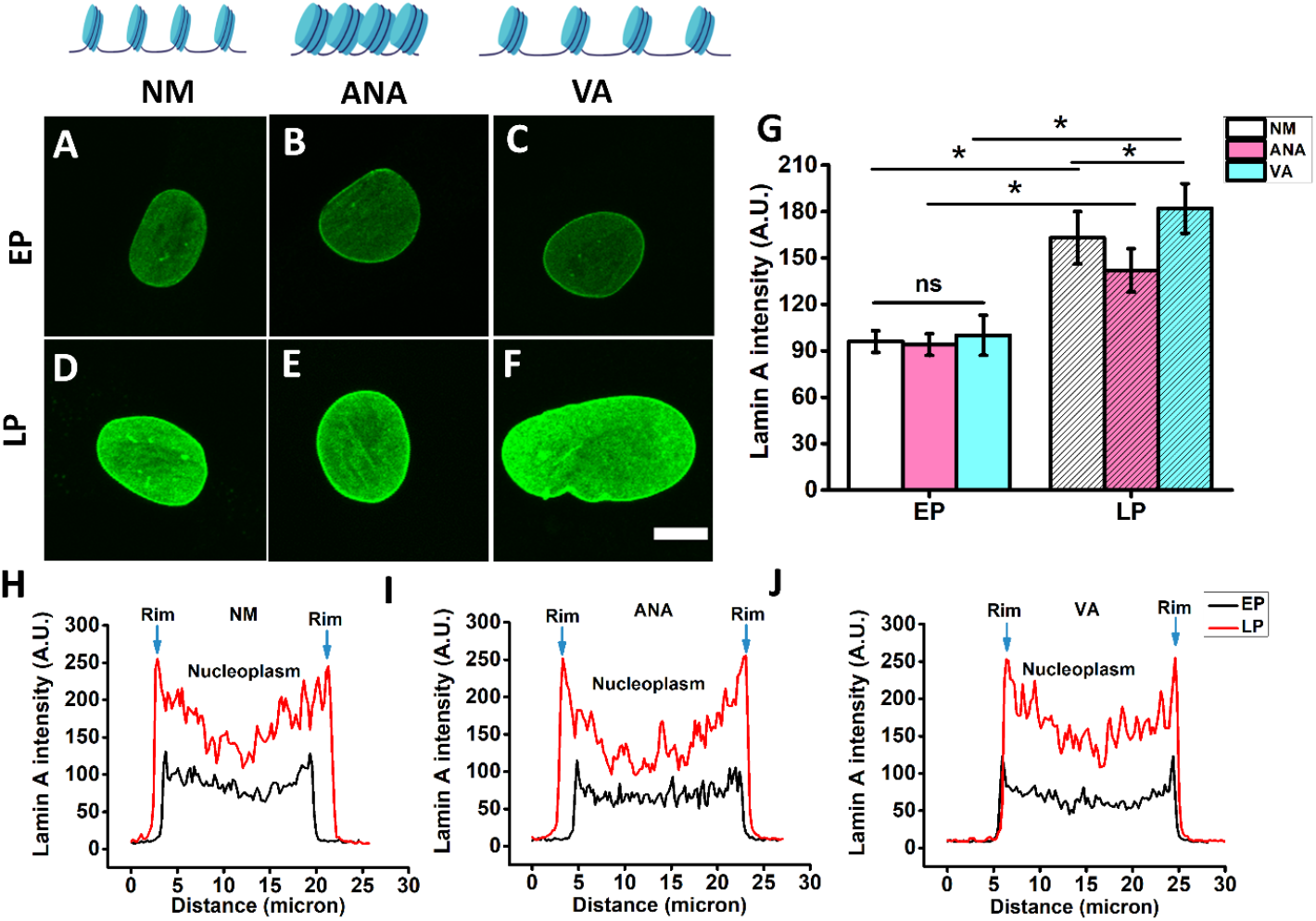
Immunofluorescence images of lamin A in early passage (EP, A-C) & late passage (LP, D-F) for hMSCs cultured in normal media (NM, A and D), with anacardic acid (ANA, B and E), and with valproic acid (VA, C and F). (G) Graph comparing lamin A intensity under various conditions (NM, ANA, VA); Distribution of lamin A intensity across nuclei in (H) normal media, (I) presence of anacardic acid and (J) presence of valproic acid (n> 60 nuclei, Scale bar = 10 µm, *p<0.05, ns = non-significant).

## Discussion

Histone acetylation/deacetylation is a critical epigenetic modification that regulates gene expression by condensing or decondensing the chromatin. While these changes happen locally for the regulation of particular genes, global histone hyperacetylation has been reported to be associated with cellular senescence (Cakouros & Gronthos, 2019; Sen et al., 2016; Sidler et al., 2017). It has been demonstrated that hyperacetylation of histones (either by inhibiting HDACs or upregulating HATs) accelerates senescence, resulting in a reduced proliferation rate. In contrast, hypoacetylation (by HATi) maintains cell proliferation (Jung et al., 2010; Lee et al., 2009). Interestingly, *in-vivo* studies with model organisms like *Drosophila, C*.*elegans*, and zebrafish have shown an opposite trend (Baell et al., 2018; Evason et al., 2008; Yu et al., 2021). Among many differences between these *in-vitro* and *in-vivo* studies, one fundamental difference is that while *in-vivo* cells experience the effect over multiple generations of cell division, most of the *in-vitro* studies are done for a single passage, with only one exception to the best of our knowledge. In the only available literature studying the effect of HDACi on long term expansion of hMSCs, Jung et al. showed that HDACi induces cellular senescence during long-term expansion through downregulation of polycomb group genes (PcGs) and upregulation of jumonji domain containing protein 3 (JMJD3) (Jung et al., 2010). The same study also demonstrated that condensing chromatin through HATi delays hMSCs senescence by upregulating polycomb genes and maintaining proliferation. Clearly, the effect of histone acetylation and deacetylation on cellular senescence in the long-term culture needs further investigation.

Consistent with study mentioned above, we found that hyperacetylation of histone with HDAC inhibitors accelerates senescence, while hypoacetylation with HAT inhibitors delays it. We confirmed this observation using well-established markers of senescence, such as the activity of beta-galactosidase and the expression of nuclear lamin-A, as well as by monitoring the population doubling time and proliferation rate. We observed that long-term passaging induced replicative senescence in all three conditions (control, HAT inhibitor ANA, and HDAC inhibitor VA), with the HAT inhibitor showing the least and the HDAC inhibitor showing the highest degree of senescence.

Interestingly, we observed a decline in hMSC proliferation in the presence of HDAC inhibitors in early passage, before any significant increase in β-galactosidase activity or lamin-A expression. As senescence is a gradual and multi-step decline in cellular physiological processes, we propose that the initial decline in proliferation is the first prominent senescence-induced outcome due to chromatin decondensation via HDAC inhibitors among various other senescence-associated phenotypes.

Another crucial issue with hMSC *in-vitro* expansion is the gradual loss of differentiation potential with serial passaging. hMSCs have tri-lineage differentiation potential, making them promising tools in tissue engineering and regenerative medicine. However, their adipogenic differentiation potential declines with long-term expansion *in-vitro* (Kureel et al., 2019b). We have shown here that when expanded in the presence of HAT inhibitors, the adipogenic differentiation potential of late-passage cells matches that of early-passage cells cultured without any inhibitor.

The findings presented in this paper introduce a fascinating possibility that warrants further investigation. Previous research from our group has demonstrated that culturing cells on softer substrates can delay senescence and preserve the differentiation potential of hMSCs and keratinocytes (Kureel et al., 2019b; Mogha et al., 2019). Moreover, multiple studies, including our own, have shown that chromatin remains in a condensed state when cells are cultured on a soft substrate, similar to the effect of HAT inhibitor treatment (Joshi et al., 2022; Killaars, Grim, Walker, Hushka, Brown, & Anseth, 2019). Given that both of these conditions - culturing cells on soft substrates and treating them with HAT inhibitors - delay senescence and maintain differentiation potential, it is conceivable that substrate stiffness influences senescence through its impact on hMSC fate across multiple passaging events, by altering histone acetylation.

In addition to other manifestations of senescence, the deterioration of nuclear morphological integrity is a critical aspect. Research has established that the nuclei of aging cells undergo various detrimental transformations such as enlarged size, loss of shape integrity, and bleb formation, which contribute to senescence (Pathak et al., 2021). While studies of short-duration drug treatment have shown that HDAC inhibitors cause nuclear abnormalities, no long-term study has been conducted to investigate the effect of histone acetylation status on nuclear structure (Kalinin et al., 2021; Stephens et al., 2018). We have shown here that HDAC inhibitors cause nuclear abnormalities, but HAT inhibitors maintain the integrity of the nuclear architecture. HAT inhibitors also protect the nucleus from DNA damage during expansion. Our results suggest that HAT inhibitors delay senescence by protecting the nucleus against DNA damage, reflected in less blebbing and maintained nuclear shape.

In summary, our study suggests that the use of HAT inhibitors can delay hMSC senescence by preserving nuclear morphology, which is evident by reduced nuclear blebbing and irregularities during long-term expansion, in addition to other reduced senescence-related phenotypes and maintained proliferation. These findings pave the way for the development of a culture system that enables the expansion of hMSCs while maintaining their self-renewal capability and differentiation potential by delaying senescence.

## Supplementary

**Figure S1.**
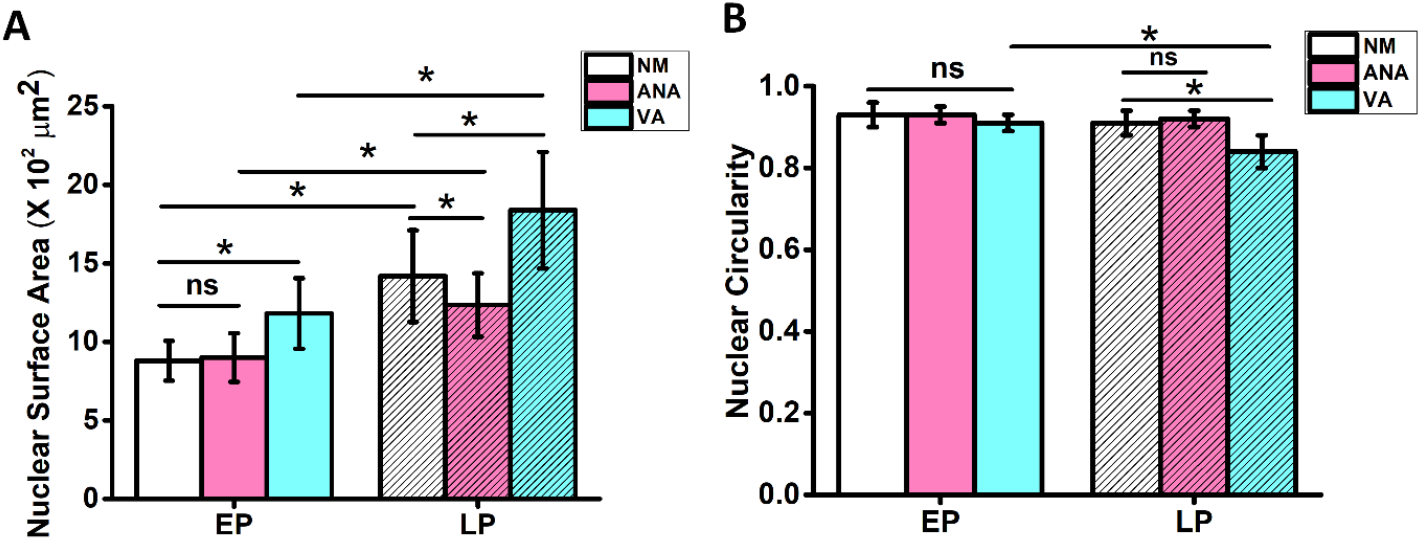
Graph comparing nuclear surface area and nuclear circularity under various conditions (NM-normal media, ANA-anacardic acid, VA-valproic acid) between early and late passage (n> 60 nuclei, Scale bar = 10 µm, *p<0.05, ns = non-significant).

